# Comprehensive evaluation of pipelines for diagnostic biomarkers of major depressive disorder using multi-site resting-state fMRI datasets

**DOI:** 10.1101/2024.03.17.584538

**Authors:** Yuji Takahara, Yuto Kashiwagi, Tomoki Tokuda, Junichiro Yoshimoto, Yuki Sakai, Ayumu Yamashita, Toshinori Yoshioka, Hidehiko Takahashi, Hiroto Mizuta, Kiyoto Kasai, Akira Kunimitsu, Naohiro Okada, Eri Itai, Hotaka Shinzato, Satoshi Yokoyama, Yoshikazu Masuda, Yuki Mitsuyama, Go Okada, Yasumasa Okamoto, Takashi Itahashi, Haruhisa Ohta, Ryu-ichiro Hashimoto, Kenichiro Harada, Hirotaka Yamagata, Toshio Matsubara, Koji Matsuo, Saori C. Tanaka, Hiroshi Imamizu, Koichi Ogawa, Sotaro Momosaki, Mitsuo Kawato, Okito Yamashita

## Abstract

The objective diagnostic and stratification biomarkers developed with resting-state functional magnetic resonance imaging (rs-fMRI) data are expected to contribute to more effective treatment for mental disorders. Unfortunately, there are currently no widely accepted biomarkers, partially due to the large variety of analysis pipelines for developing them. In this study we comprehensively evaluated analysis pipelines using a large-scale, multi-site fMRI dataset for major depressive disorder (MDD) (1162 participants from eight imaging sites). We explored the combinations of options in four subprocesses of analysis pipelines: six types of brain parcellation, four types of estimations of functional connectivity (FC), three types of site difference harmonization, and five types of machine learning methods. 360 different MDD diagnostic biomarkers were constructed using the SRPBS dataset acquired with unified protocols (713 participants from four imaging sites) as a discovery dataset and evaluated with datasets from other projects acquired with heterogeneous protocols (449 participants from four imaging sites) for independent validation. To identify the optimal options regardless of the discovery dataset, we repeated the same procedure after swapping the roles of the two datasets. We found pipelines that included Glasser’s parcellation, tangent-covariance, no harmonization, and non-sparse machine learning methods tended to result in high classification performance. The diagnosis results of the top 10 biomarkers showed high similarity, and weight similarity was also observed between eight of the biomarkers, except two that used both data-driven parcellation and FC computation. We applied the top 10 pipelines to the datasets of other mental disorders (autism spectral disorder: ASD and schizophrenia: SCZ) and eight of the ten biomarkers showed sufficient classification performances for both disorders, except two pipelines that included Pearson correlation, ComBat harmonization and random forest classifier combination.

**Highlights:**

- We evaluated the analysis pipelines of rsFC biomarker development.
- Four subprocesses in them were investigated with two multi-site datasets.
- Glasser’s parcellation, tangent covariance, and non-sparse methods were preferred.
- The weight patterns of eight of the top 10 biomarkers showed high commonality.
- Eight of the top 10 pipelines were successful for developing SCZ/ASD biomarkers.

## 1. Introduction

Most psychiatric disorders, including major depressive disorder (MDD), are diagnosed based on the Diagnostic & Statistical Manual of Mental Disorders, such as DSM-5 or the International Classification of Diseases (ICD). Their diagnosis is based on visible symptoms and medical interviews. Although the overlapping phenotypes among multiple psychiatric disorders complicates selecting appropriate treatment at an early stage, no objective, practical marker yet leads to more refined diagnoses and targeted treatment (Arnow et al., 2015; Kapur et al., 2012).

Non-invasive brain imaging techniques are expected to elucidate the patterns of brain structure and function, that is, neurophenotypes, which characterize these disorders (Craddock et al., 2013; van Essen and Ugurbil, 2012). In particular, acquiring a resting-state fMRI (rs-fMRI) is easy and can be used to develop classification markers between healthy and psychiatric or developmental disorder populations (Clare Kelly et al., 2008; van Essen and Ugurbil, 2012), such as Alzheimer’s disease (Chen et al., 2011; Greicius et al., 2004), schizophrenia (Calhoun et al., 2012; Garrity et al., 2007; Jafri et al., 2008; Yoshihara et al., 2020; Zhou et al., 2007), autism spectrum disorder (Plitt et al., 2015; Yahata et al., 2016), depression (Craddock et al., 2009; Ichikawa et al., 2020; Yamashita et al., 2020), and attention-deficit hyperactivity disorder (Milham et al., 2012). In recent years, several worldwide projects have acquired large-scale, rs-fMRI data (Koike et al., 2021; Martino et al., 2013; Tanaka et al., 2021; van Essen et al., 2013), and the understanding of psychiatric disorders and the development of markers is progressing. Unfortunately, no practical diagnostic markers have been identified yet (Castellanos et al., 2013; Kapur et al., 2012), perhaps explained by two factors: the absence of optimal pipelines and the lack of generalization capability.

The development of classification biomarkers using rs-fMRI is comprised of several processes, and there are multiple methods in every process as well as innumerable pipelines (Arbabshirani et al., 2017; Brown and Hamarneh, 2016; Wolfers et al., 2015). The diversity of pipelines has a sizable effect on the diagnostic and generalization performance, and only a few reports have searched for the best ones (Dadi et al., 2019; Mellema et al., 2022; Pervaiz et al., 2020). The absence of standard pipelines reduces the reliability of rs-fMRI biomarkers. Since no widely accepted standard pipeline has been established for optimal biomarker development (Carp, 2012), the identification of a suitable pipeline in the field of psychiatric disorders is critical to develop diagnostic markers for rs-fMRI.

Since the majority of older biomarkers were made based on single-site data to avoid unknown effects of the heterogeneity of discovery datasets, they often exhibit a lack of generalization capability. The differences of scanners, imaging procedures, and instructions to participants might also affect rs-fMRI data. To develop markers with high generalization capability, discovery datasets must be comprised of large-size, multi-site data; such methods must be devised. Cross-validated modeling is an effective approach to avoid overfitting. After developing a marker using a multi-site dataset, it must be validated using an independent multi-site dataset. This validation test enhances the marker’s reliability. Several methods have minimized the inter-site differences of rs-fMRI, including ComBat (Fortin et al., 2018, 2017; Johnson et al., 2007; Yu et al., 2018) and traveling-subject harmonization (Yamashita et al., 2019). These methodological approaches are also expected to enhance the generalization capability of markers.

Our aim in this study is to explore an optimal pipeline to develop an MDD diagnostic biomarkers with high classification performance for an independent validation dataset using a large-size, multi-site rs-fMRI dataset. In this study, we also verified the effectiveness of the state-of-the-art methods (Glasser’s surface-based parcellation and distance correlation) and a site-difference harmonization method, which were not compared in previous exploratory studies, and prepared an independent dataset for the validation of a marker’s generalization capability. By swapping the roles of the discovery and validation datasets, we also confirmed that a pipeline’s effectiveness did not depend on the discovery dataset.

## 2. Materials and methods

### Ethics statement

Every participant in all the datasets provided written informed consent. All the recruitment procedures and experimental protocols were conducted in accordance with the Declaration of Helsinki and approved by the institutional review boards of the principal investigators’ respective institutions (Advanced Telecommunications Research Institute International [approval numbers: 13-133, 14-133, 15-133, 16-133, 17-133, and 18-133], Hiroshima University [E-38], Kyoto Prefectural University of Medicine [RBMR-C-1098], Showa University [SWA] [B-2014-019 and UMIN000016134], the University of Tokyo [UTO] Faculty of Medicine [3150], Kyoto University [C809 and R0027], and Yamaguchi University [H23-153 and H25-85]).

### 2.1. Patients and subjects

We analyzed the same datasets previously presented in Yamashita et al., 2020: (1) Dataset I contained data from 713 participants (564 healthy controls (HCs) from four sites and 149 MDDs from three sites); (2) Dataset II contained data from 449 participants (264 HCs and 185 MDDs from four independent sites); (3) Dataset III contained data from 231 participants (125 with autism spectrum disorder (ASD) from two sites and 106 with schizophrenia (SCZ) from three sites). For more details on the dataset, see Tables 1 and 2 and Supplemental Table 1.

**Table 1.**
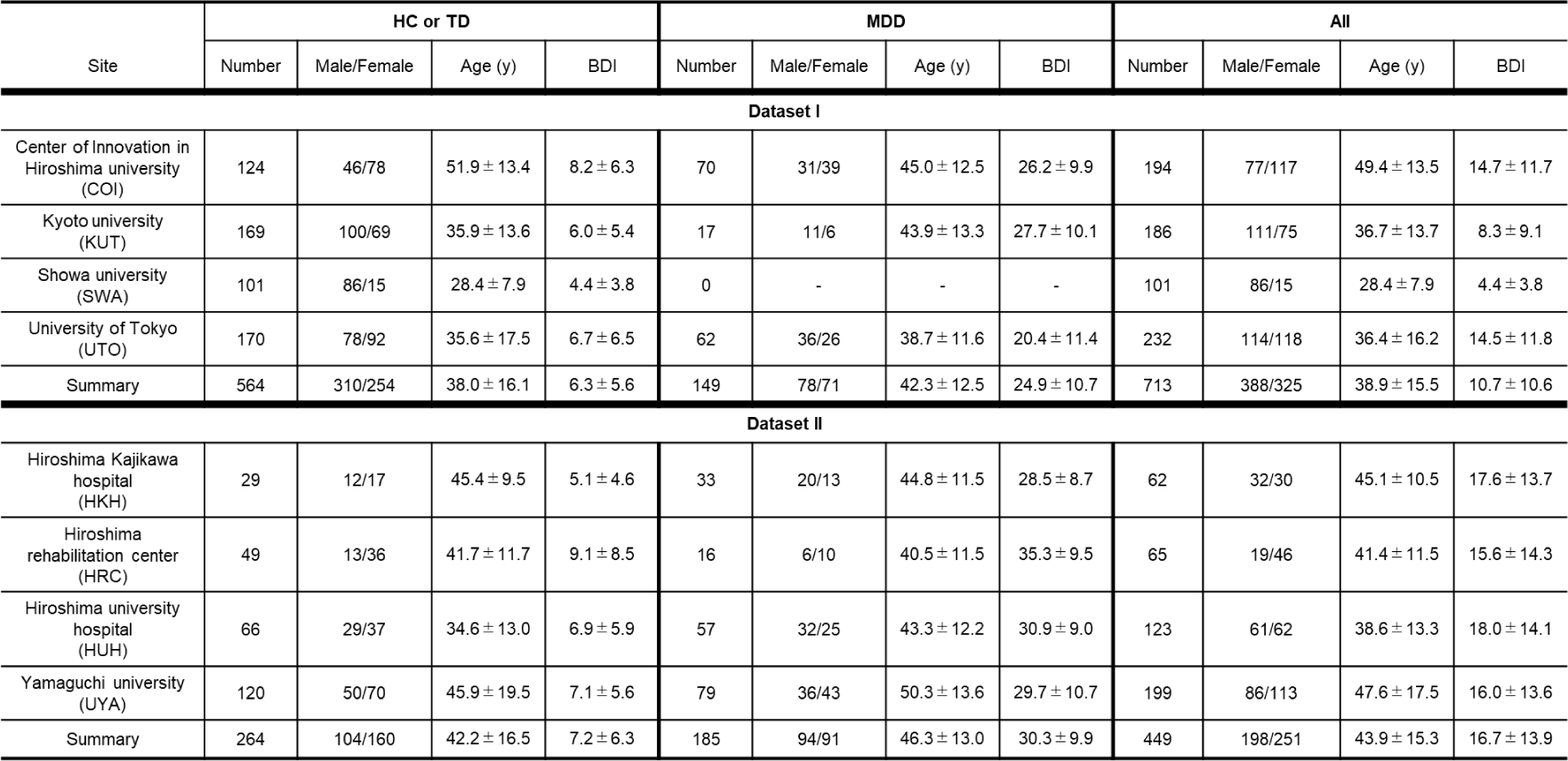
Demographic information of participants: Dataset I contained data from 713 participants (564 HCs from four sites and 149 MDDs from three sites). All data were acquired using a unified imaging protocol (SRPBS DecNef project). Dataset II contained data from 449 participants (264 HCs and 185 MDDs) from completely different four sites from four sites of dataset I. See Supplemental Table 1 for more detailed information protocols.

| Site | HC or TD |  |  |  | MDD |  |  |  | All |  |  |  |
| --- | --- | --- | --- | --- | --- | --- | --- | --- | --- | --- | --- | --- |
|  | Number | Male/Female | Age (y) | BDI | Number | Male/Female | Age (y) | BDI | Number | Male/Female | Age (y) | BDI |
| Dataset I |  |  |  |  |  |  |  |  |  |  |  |  |
| Center of Innovation in Hiroshima university (COI) | 124 | 46/78 | 51.9 ± 13.4 | 8.2 ± 6.3 | 70 | 31/39 | 45.0 ± 12.5 | 26.2 ± 9.9 | 194 | 77/117 | 49.4 ± 13.5 | 14.7 ± 11.7 |
| Kyoto university (KUT) | 169 | 100/69 | 35.9 ± 13.6 | 6.0 ± 5.4 | 17 | 11/6 | 43.9 ± 13.3 | 27.7 ± 10.1 | 186 | 111/75 | 36.7 ± 13.7 | 8.3 ± 9.1 |
| Showa university (SWA) | 101 | 86/15 | 28.4 ± 7.9 | 4.4 ± 3.8 | 0 | - | - | - | 101 | 86/15 | 28.4 ± 7.9 | 4.4 ± 3.8 |
| University of Tokyo (UTO) | 170 | 78/92 | 35.6 ± 17.5 | 6.7 ± 6.5 | 62 | 36/26 | 38.7 ± 11.6 | 20.4 ± 11.4 | 232 | 114/118 | 36.4 ± 16.2 | 14.5 ± 11.8 |
| Summary | 564 | 310/254 | 38.0 ± 16.1 | 6.3 ± 5.6 | 149 | 78/71 | 42.3 ± 12.5 | 24.9 ± 10.7 | 713 | 388/325 | 38.9 ± 15.5 | 10.7 ± 10.6 |
| Dataset II |  |  |  |  |  |  |  |  |  |  |  |  |
| Hiroshima Kaijika hospital (HKH) | 29 | 12/17 | 45.4 ± 9.5 | 5.1 ± 4.6 | 33 | 20/13 | 44.8 ± 11.5 | 28.5 ± 8.7 | 62 | 32/30 | 45.1 ± 10.5 | 17.6 ± 13.7 |
| Hiroshima rehabilitation center (HRC) | 49 | 13/36 | 41.7 ± 11.7 | 9.1 ± 8.5 | 16 | 6/10 | 40.5 ± 11.5 | 35.3 ± 9.5 | 65 | 19/46 | 41.4 ± 11.5 | 15.6 ± 14.3 |
| Hiroshima university hospital (HUH) | 66 | 29/37 | 34.6 ± 13.0 | 6.9 ± 5.9 | 57 | 32/25 | 43.3 ± 12.2 | 30.9 ± 9.0 | 123 | 61/62 | 38.6 ± 13.3 | 18.0 ± 14.1 |
| Yamaguchi university (UYA) | 120 | 50/70 | 45.9 ± 19.5 | 7.1 ± 5.6 | 79 | 36/43 | 50.3 ± 13.6 | 29.7 ± 10.7 | 199 | 86/113 | 47.6 ± 17.5 | 16.0 ± 13.6 |
| Summary | 264 | 104/160 | 42.2 ± 16.5 | 7.2 ± 6.3 | 185 | 94/91 | 46.3 ± 13.0 | 30.3 ± 9.9 | 449 | 198/251 | 43.9 ± 15.3 | 16.7 ± 13.9 |

**Table 2.** Demographic characteristics of participants for other disorders: In development of ASD & SCZ diagnostic marker, rs-fMRI data acquired from 125 ASDs (two sites) and 106 SCZs (three sites) was used. For TD and HC, the same HC as dataset I was used. All data were acquired using a unified imaging protocol (SRPBS DecNef project).

| Dataset III |  |  |  |  |  |  |
| --- | --- | --- | --- | --- | --- | --- |
| Site | ASD |  |  | SCZ |  |  |
|  | Number | Male/Female | Age (y) | Number | Male/Female | Age (y) |
| Center of Innovation in Hiroshima university (COI) | 0 | - | - | 0 | - | - |
| Kyoto university (KUT) | 0 | - | - | 51 | 26/25 | 41.0 ± 10.7 |
| Showa university (SWA) | 115 | 100/15 | 32.1 ± 7.8 | 19 | 15/4 | 42.9 ± 8.4 |
| University of Tokyo (UTO) | 10 | 9/1 | 37.0 ± 9.6 | 36 | 24/12 | 31.4 ± 10.3 |
| Summary | 125 | 109/16 | 32.5 ± 8.0 | 106 | 65/41 | 38.1 ± 11.2 |

### 2.2. Preprocessing

We preprocessed the rs-fMRI data using fMRIPrep version 1.0.8 (Esteban et al., 2019). The first 10 s of the data were discarded to allow for T1 equilibration. The following are the preprocessing steps: slice-timing correction, realignment, coregistration, distortion correction using a field map, segmentation of the T1-weighted structural images, normalization to Montreal Neurological Institute (MNI) space, and spatial smoothing with an isotropic Gaussian kernel of 6-mm full width at half maximum. “Fieldmap-less” distortion correction was performed for dataset II due to a lack of field-map data. For more details on the pipelines, see http://fmriprep.readthedocs.io/en/1.0.8/workflows.html. For the data of six participants in dataset II, coregistration was unsuccessful, and so we excluded them from further analysis.

### 2.3. Parcellation

We considered six methods for extracting the brain’s regions of interest (ROIs): five pre-defined parcellations, which were actually used for the development of depression diagnosis and stratification biomarkers (Ichikawa et al., 2020; Tokuda et al., 2018; Yamashita et al., 2020; Yoo et al., 2018), and one data-driven parcellation that was reported as the best choice (Dadi et al., 2019, Figure 1).

**Figure 1.**
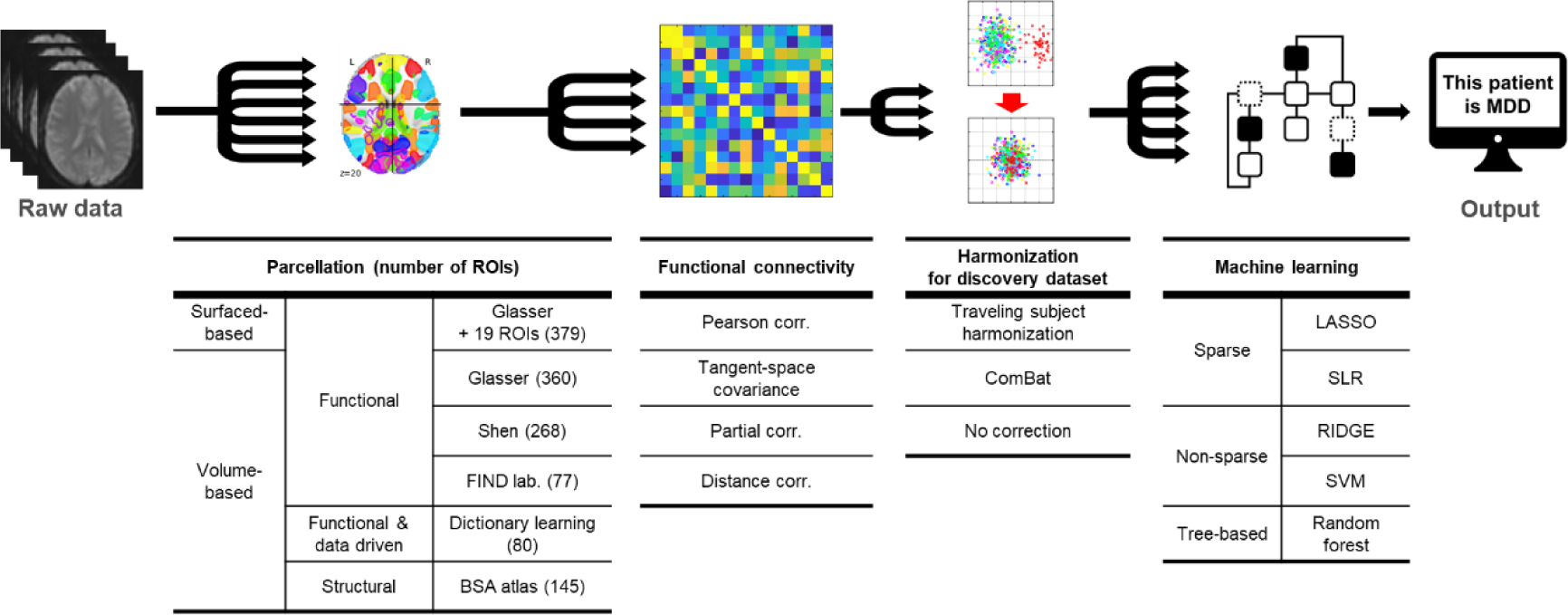
Design for comprehensive exploration of analysis pipelines of rsFC MDD biomarker: After preprocessing with fmriprep-ciftify, 360 markers were constructed using 360 pipelines (6 parcellations × 4 functional connectivity × 3 site-bias harmonization × 5 machine learning).

#### Pre-defined parcellation

1. Glasser’s surface-based method: 379 ROIs, including 360 cortical parcels and 19 subcortical parcels (Glasser et al., 2016), utilized by ciftify toolbox version 2.0.2-2.0.3 (Dickie et al., 2019).
2. Glasser’s volume-based method [https://figshare.com/articles/dataset/HCP-MMP1_0_projected_on_fsaverage/3498446], which has only 360 cerebral cortex ROIs.
3. Shen’s atlas, derived on a group-wise spectral clustering algorithm, which has 268 ROIs (Shen et al., 2013).
4. Brainvisa, anatomically defined in the Brainvisa Sulci Atlas (BSA atlas), which has anatomically-defined 145 ROIs covering the entire cerebral cortex [http://brainvisa. Info; Perrot et al., 2011].
5. FIND lab’s parcellation (Shirer et al., 2012), based on functional (rather than structural) ROIs from which we selected 78 ROIs, excluding cerebellum-related ROIs **Data-driven parcellation:** We defined ROIs using a linear decomposition method: dictionary learning (Mensch et al., 2016). A data-driven atlas was constructed using the discovery dataset. We fixed the number of components to dim = 80, which is the optimal number (Dadi et al., 2019).

### 2.4. Functional connectivity (FC) matrix

We considered four FC calculation methods (Figure 1): Pearson’s full correlation coefficient, tangent-space covariance (Varoquaux et al., 2010), partial correlation, and distance correlation (Yoo et al., 2019). For each participant, the FC was calculated from rs-fMRI BOLD signals across the ROIs for each parcellation. All the FCs were calculated using Nilearn Python library version 0.6.1 [https://nilearn.github.io/stable/index.html]. We used the FC values of the lower triangular matrix of the connectivity matrix and applied Fisher’s z-transformation to each FC, except for the tangent-space covariance. We calculated the tangent-space covariances both of the discovery and validation datasets, based on the group mean values of the former.

### 2.5. Preprocessing for ROI time series

Physiological noise regressors were extracted by applying CompCor (Behzadi et al., 2007). To remove several sources of spurious variance, we used a linear regression with 12 regression parameters: six motion parameters, average signals over the whole brain, and five anatomical CompCor components.

We applied a temporal bandpass filter to the time series using a second-order Butterworth filter with a pass band between 0.01 and 0.08 Hz to restrict the analysis to low-frequency fluctuations, which are characteristic of rs-fMRI BOLD activity (Ciric et al., 2017).

A scrubbing process based on head motion was conducted using frame displacement (FD, Power et al., 2014), which was calculated using Nipype (https://nipype.readthedocs.io/en/latest/). We removed the volumes with FD > 0.5 mm, unlike a previous study (Power et al., 2014). Using this threshold, 6.3% ± 13.5 volumes (mean ± SD) were removed per rs-fMRI session from all the datasets. Subjects whose ratio of excluded volumes by scrubbing exceeded the mean + 3 SD were removed, resulting in 48 participants being removed from all the datasets. Thus, we included 683 participants (545 HCs and 138 MDDs) in dataset I, 444 participants (263 HCs and 181 MDDs) in dataset II, and 218 participants (116 ASDs and 102 SCZs) in dataset III.

### 2.6. Harmonization

We utilized three options as site difference harmonization methods in the discovery dataset (Figure 1): traveling-subject harmonization, ComBat, and no correction.

Traveling-subject harmonization (TS method) enables us to estimate measurement bias (**m**) by controlling participant bias (**p**) using the traveling-subject dataset (Supplemental Table 2), in which multiple participants traveled to multiple recording sites that recorded their rs-fMRIs with identical recording protocols (Yamashita et al., 2019). For each connectivity, the regression model can be written:

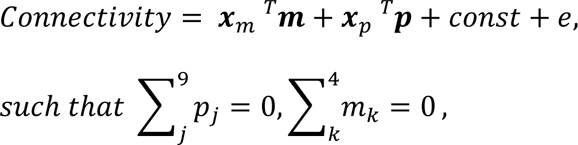

where **m** represents the measurement bias (4 sites × 1), **p** represents the participant factor (nine traveling subjects × 1), *const* represents the average FC values across all the participants from all sites, and *e* ∼ *N*(0, γ^−1^) represents noise. **x**m and **x**p are vectors represented by 1-of-K binary coding. **x**m for measurement bias ***m*** belonging to site *k* is a binary vector where all the elements equal zero, except for element *k*, which equals 1. Measurement biases were removed by subtracting the estimated measurement biases. Thus, the harmonized FC values were set:

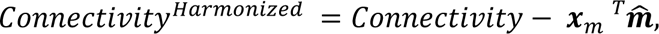

which represents the estimated measurement bias. More detailed information was previously described (Yamashita et al., 2019). ComBat: the ComBat harmonization method (Dansereau et al., 2017; Fortin et al., 2018, 2017; Johnson et al., 2007) 2007) is a well-known control for site differences in FC. Although the TS method requires that the traveling-subject dataset is acquired in advance, the ComBat approach allows site differences to be corrected using only discovery data. We performed ComBat harmonization to correct only for site differences while keeping fc correlated with two biological covariates, age and sex. No correction: The deleterious effect of site differences for prediction accuracy decreased as the total sample size increased (Dansereau et al., 2017). We did not apply harmonization to the validation dataset to determine the effects of harmonization on biomarker construction.

### 2.7. Machine learning

We considered five supervised machine learning methods (Figure 1): *1)* least absolute shrinkage and selection operator (LASSO) was performed using the *“lassoglm”* function, and we set *“NumLambda”* to 25 and *“CV”* to 10. λ was determined according to the one standard error rule in which we selected the largest λ within the standard deviation of the minimum prediction error. *2)* Sparse Logistic Regression (SLR) was performed using the *“biclsfy_slrvar”* function, and we set *“nlearn”* to 300 and *“usebias”* to 1 (https://bicr.atr.jp/~oyamashi/SLR_WEB.html, Yamashita et al., 2008). *3)* RIDGE was performed using the *“fitclinear”* function, and we set *“Solver”* to {*“sgd”*, *“lbfgs”*}, *“OtimizeHyperparameters”* to *“Lambda”* and *“Kfold”* to 10. *4)* Support Vector Machine (SVM) was performed using the *“fitcsvm”* function, and we set *“KernelScale”* and *“Boxconstraint”* to the optimization parameters derived from the *“bayesopt”* function. *5)* Random Forest was performed using the *“TreeBagger”* function, and we set *“NumTrees”* to 1000 and *“PredictorSelection”* to *“interaction-curvature”*. All the functions described here were executed in MATLAB (R2018b, Mathworks, USA).

### 2.8. Constructing and validating a MDD classifier

We constructed a diagnostic brain network biomarker for MDD that classified between HCs and MDDs using the discovery dataset, e.g., dataset I, based on each FC value (Supplemental Figure 1). To avoid overfitting issues, we used the 10-fold nested cross-validation (CV) method, which is based on a previously proposed method (Yamashita et al., 2020) with a slight modification to the standardization of the validation datasets. We first divided the whole dataset I into a training set (9 of 10 folds), which was used for training a model, and a test set (1 of 10 folds), which was used for testing it. To prevent bias due to the differences in the numbers of the two groups, we used an undersampling method to equalize the numbers between the MDD and HC groups (Wallace et al., 2011). Since only a subset of the training set is used after undersampling, we repeated the random sampling procedure ten times (i.e., subsampling). When we performed the undersampling, we matched the mean ages between the MDD and HC groups in each undersampling and standardized both the undersampled training subset and the test set with the mean and the standard deviation values of each FC in each undersampling. We then fit a model to each subsample and created ten classifiers. The mean classifier-output value (diagnostic probability) was indicative of the classifier output. Subjects with a diagnostic probability greater than 0.5 were considered MDD patients.

We tested the generalization capability of the diagnostic markers using the validation dataset, e.g., dataset II. Then the diagnostic probability of each subject was calculated after the FC was standardized by the mean value and the standard deviation of the training subset.

### 2.9. Evaluation criteria

We calculated the area under the curve (AUC), the accuracy, the sensitivity, the specificity, and the Matthews correlation coefficient (MCC, Chicco, 2017).

Furthermore, since using only AUC may not accurately compare the classification performance for the imbalanced data, we calculated a “composite score” that combined AUC and the scale-adjusted MCC:

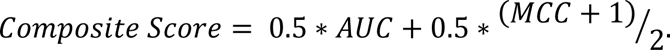

We also quantified the instability of the classification results between the 10-fold CV result for the discovery dataset and the application result for the validation dataset and defined “instability”:

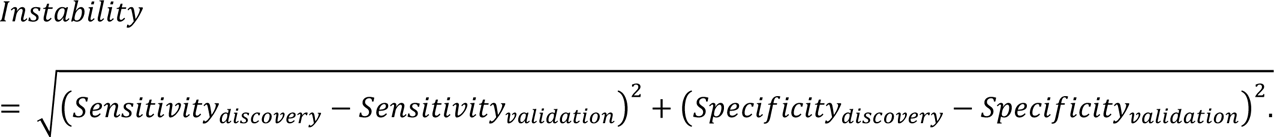

### 2.10. Confirmation of pipeline stability

We confirmed the pipeline stability by dataset-role swapping. We constructed MDD diagnostic biomarkers using identical pipelines, except for the traveling-subject harmonization method when dataset II was used as the discovery dataset and dataset I as the validation dataset. We calculated their composite scores, instability, and diagnostic probabilities on each marker. The correlation coefficient was calculated of the marker’s diagnostic probabilities before and after dataset-role swapping.

### 2.11. Marker and pipeline ranking

For the 240 pipelines to which both datasets can be applied as discovery datasets, we standardized the composite scores and instability values before and after the dataset-role swapping and the correlation coefficient values of the diagnostic probability. The final pipeline ranking was determined by the sum of these five standardized values (Table 5).

### 2.12. Evaluation of similarity of important functional connectivity

We examined the network-level similarity of the important functional connectivity among the top 10 highest diagnostic performance MDD biomarkers. Since each diagnostic marker has 100 classifiers, the contribution of each FC to the diagnostic marker can be quantified by the sum of the absolute values of the weight values or the out-of-bag predictive importance. To compare the similarity of FC patterns between markers constructed using different parcellation options, we applied the FCs extracted as the top 5% of high contribution FCs in each marker to the same atlas of seven brain networks (Buckner et al., 2011; Choi et al., 2012; Thomas Yeo et al., 2011). The ROIs that comprised each FC were counted where they were classified on each brain network label and divided by twice the total number of the top 5% high contribution FCs. Since biases exist in the number of ROIs classified into each network depending on the parcellation, we calculated the existence probability of the ROIs of all the FCs on the brain network for each parcellation and divided the existence probability of the top 5% FCs by the existence probability of the whole FCs. The correlation coefficient of the brain network utilization between any two markers’ high contribution FCs was used as a quantitative index of marker similarity. The similarity is important if the correlation coefficient among two different utilization rates was significantly higher than a randomness threshold. We randomly selected an arbitrary 5% of the FCs from each parcellation and calculated the corrected utilization rate on Yeo’s brain network and the correlation coefficient between these two utilization rates. We conducted this operation 10,000 times and set a statistical significance at a certain threshold (permutation test, P < 0.05, 1-sided).

### 2.13. Evaluation of pipelines for other disorders

Using the top 10 pipelines suitable for constructing diagnostic markers for MDD, we confirmed the applicability of those pipelines to other two disorders: schizophrenia (SCZ) and autism spectrum disorders (ASD). We developed diagnostic biomarkers for SCZ and ASD using the same process as for the MDD diagnostic biomarkers. The same data as the HCs of Section 2.1 were used for the HC and typical development (TD) subjects, and all the data were collected in the DecNef Project Brain Data Repository (https://bicr.atr.jp/decnefpro/data; Tables 1 and 2, 564 HC or TD, 106 SCZ, and 125 ASD patients).

## 3. Results

### 3.1. Comparison of diagnostic performances with all markers

Focusing on the four steps of the whole marker construction processes (Figure 1), we searched for the optimal pipeline for the construction of MDD diagnostic biomarkers with superior classification performance for both the discovery dataset and the validation dataset (Supplemental Figure 1). We utilized 360 different pipelines and constructed 360 different brain network markers for MDD and distinguished between HCs and MDDs using dataset I as a discovery dataset. Then we applied all the markers in the validation dataset (dataset II). For each one, we obtained the area under the curve (AUC) and the Matthews correlation coefficient (MCC), and the sensitivity and specificity were obtained for the discovery and validation datasets. Furthermore, we calculated “composite scores” by combining the AUCs and the scale-adjusted MCCs to compare the diagnostic performances and the “instability” index, which is the distance between two sensitivities and two specificities, to evaluate the stability of the discrimination result between the discovery and validation datasets.

According to the composite scores, we first studied the importance of the choice of each development process in the marker pipelines: the choice of parcellations, functional connectivity, harmonization, and machine learning (Figure 1). In the parcellation process (Figure 2A), Glasser + 19 ROIs (379) and dictionary learning (80) showed higher composite scores for both dataset I (discovery) and dataset II (validation). In the functional connectivity process (Figure 2B), Pearson’s full correlation and tangent-space covariance outperformed the rest. In the harmonization process (Figure 2C), some markers using pipelines with the traveling-subject harmonization approach scored very high, although those using traveling-subject pipelines generally scored slightly lower than markers with pipelines that did not apply any inter-site correction. On the other hand, the diagnostic performances of the markers in the discovery dataset using a pipeline containing the ComBat option were significantly lower than other markers. Last, in the machine learning process (Figure 2D), the non-sparse methods, RIDGE and SVM, showed higher composite scores, and some of the markers constructed using the pipeline, including the random forest method, showed very high diagnostic performance for dataset II.

**Figure 2.**
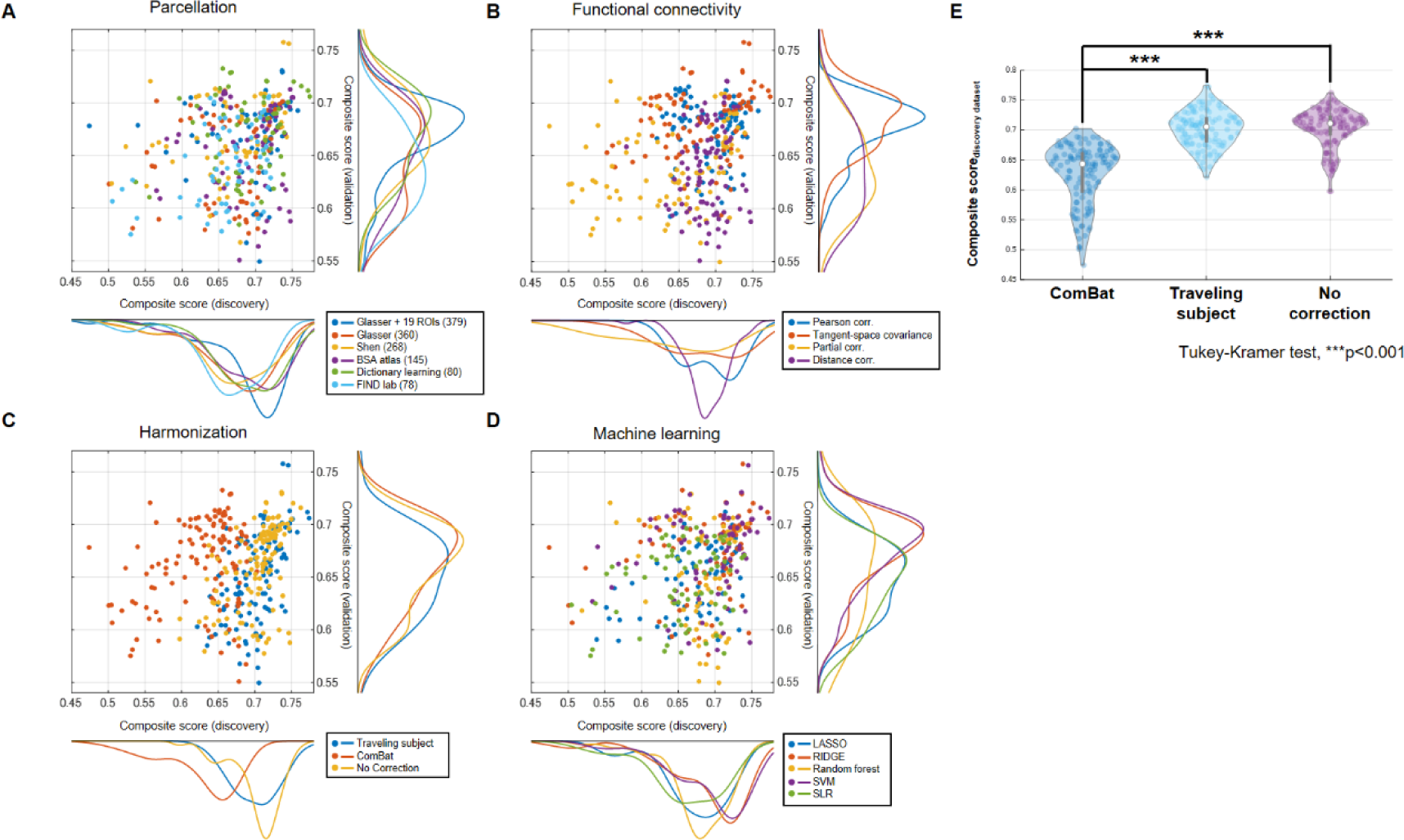
Classification performances of markers constructed with 360 pipelines: *A-D)* Distribution of composite scores both in discovery and validation datasets for each process. *A)* Parcellation, *B)* Functional connectivity estimation, *C)* Site-bias harmonization, *D)* Machine learning. *E)* Distribution of composite scores of all diagnostic markers for each harmonization method in discovery dataset. Composite scores of markers constructed using ComBat show significant lower values than those of remaining markers (Tukey-Kramer test, ***p<0.001).

Since significant differences were identified between the options in the composite score of dataset I (discovery) in the inter-site harmonization process (Figure 2E), we investigated with markers that were constructed using only 240 pipelines that did not use ComBat as a harmonization option in the following analysis of effective methods in each development process. We considered the optimum method for each development process by comparing the composite scores of the markers of all the pipelines, which can be determined when the option in a certain process is fixed to one choice. We used a multiway analysis of variance to evaluate the effects of multiple factors on the averaged composite scores using the ANOVAN function in MATLAB (Table 3, MathWorks, R2018b). The averaged composite score was the mean value of the composite scores of both datasets and reflected the marker’s generalization capability. In this multiple comparison for a four-way ANOVA, the averaged composite scores were standardized. Except for the parcellation and harmonization combination, all four main terms and five of the six interaction terms seemed to significantly affect the averaged composite score. Regarding parcellation, surface-based parcellation (Glasser + 19 ROIs (379)) and data-driven parcellation (dictionary learning (80)) led to maximal performance (Figure 3A). Tangent-space covariance and Pearson’s full correlation as a functional connectivity tended to outperform the partial and distance correlations. Indeed, they performed better on average (Figure 3B). Regarding the harmonization approach, we found no significant difference between the no-correction option and traveling-subject harmonization (Figure 3C). As a machine learning method, the result shows that non-sparse linear classifiers, including support vector machine (SVM) and RIDGE, outperformed the other approaches (Figure 3D). Based on this analysis, the recommended methods for each process are summarized in Table 4.

**Figure 3.**
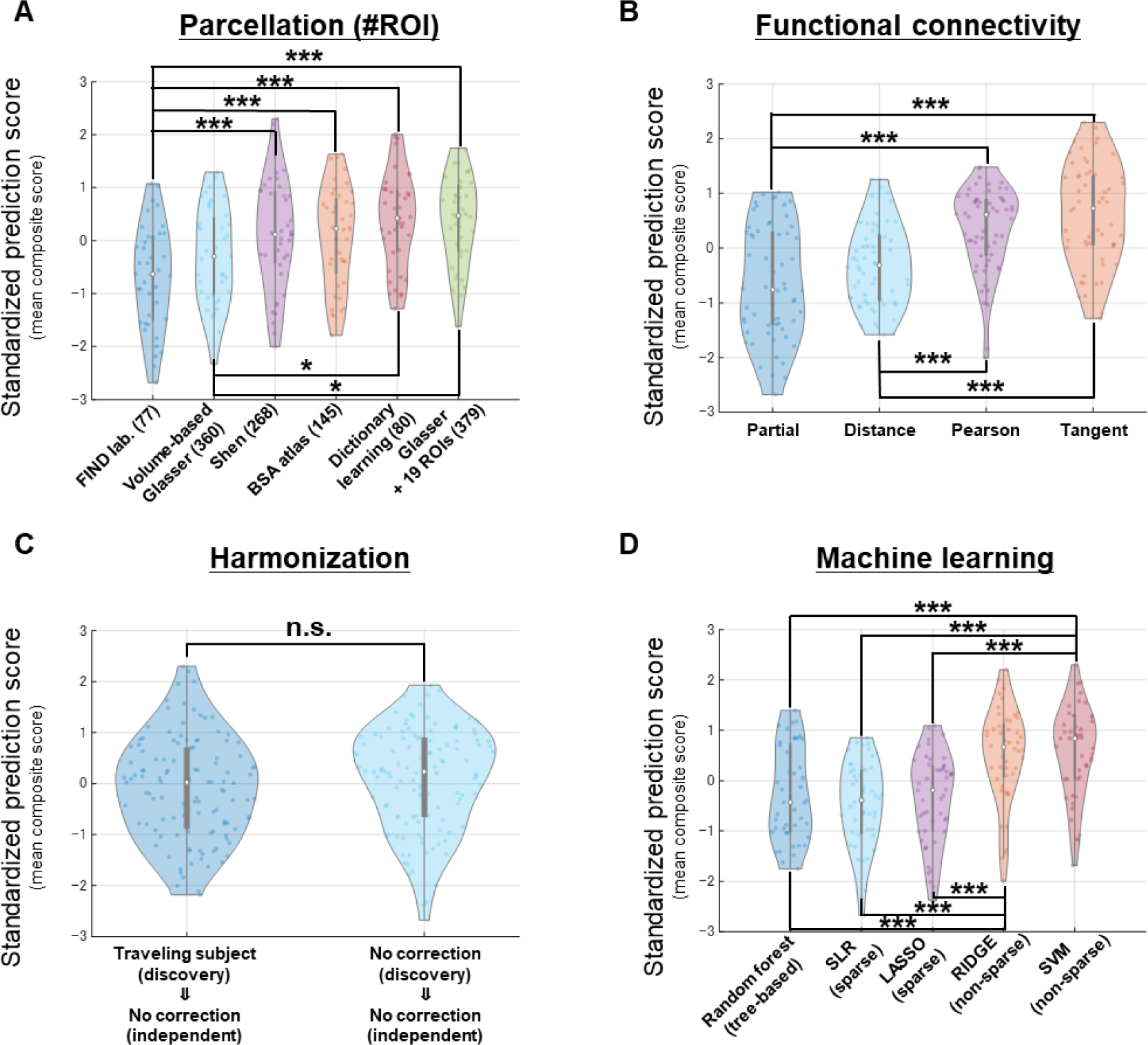
Impact of each method choice on prediction accuracy in each development process: Distribution of diagnostic performances for each method selected in each development process when excluding markers constructed using ComBat method. *A)* In Parcellation process, Glasser + 19 ROIs (379) and dictionary learning (80) performed better. *B)* In Functional connectivity process, Pearson’s full correlation and tangent-space covariance performed better. *C)* In Harmonization process, there is no significant difference. *D*) In Machine learning process, non-sparse methods outperformed sparse methods. In this analysis, standardized prediction scores are used by standardization of averaged composite scores of both datasets (Tukey-Kramer test, ***p<0.001, **p<0.01, *p<0.05).u

**Table 3.** Multi-way ANOVA for mean composite score except markers using ComBat: After eliminating markers constructed using ComBat options, four-way ANOVA was conducted on averaged composite scores of all markers, focusing on four development processes.

| Source | Sum sq. | df | Mean sq. | F | Sig. |
| --- | --- | --- | --- | --- | --- |
| Parcellation | 0.03345 | 5 | 0.00669 | 37.75 | 1.21*10<sup>-25</sup> |
| Functional connectivity (FC) | 0.05469 | 3 | 0.01823 | 102.86 | 1.70*10<sup>-37</sup> |
| Harmonization | 0.00194 | 1 | 0.00194 | 10.95 | 0.00115 |
| Machine learning | 0.04784 | 4 | 0.01196 | 67.49 | 1.16*10<sup>-33</sup> |
| Parcellation<br>× FC | 0.02011 | 15 | 0.00134 | 7.57 | 1.23*10<sup>-12</sup> |
| Parcellation<br>× Harmonization | 0.00171 | 5 | 0.00034 | 1.93 | 0.09208 |
| Parcellation<br>× Machine learning | 0.01152 | 20 | 0.00058 | 3.25 | 1.64*10<sup>-5</sup> |
| FC<br>× Harmonization | 0.00452 | 3 | 0.00151 | 8.5 | 2.79*10<sup>-5</sup> |
| FC<br>× Machine learning | 0.01435 | 12 | 0.0012 | 6.75 | 8.41*10<sup>-10</sup> |
| Harmonization<br>× Machine learning | 0.01054 | 4 | 0.00263 | 14.87 | 2.18*10<sup>-10</sup> |
| Error | 0.02906 | 164 | 0.00018 |  |  |
| Total | 0.23324 | 236 |  |  |  |

**Table 4.**
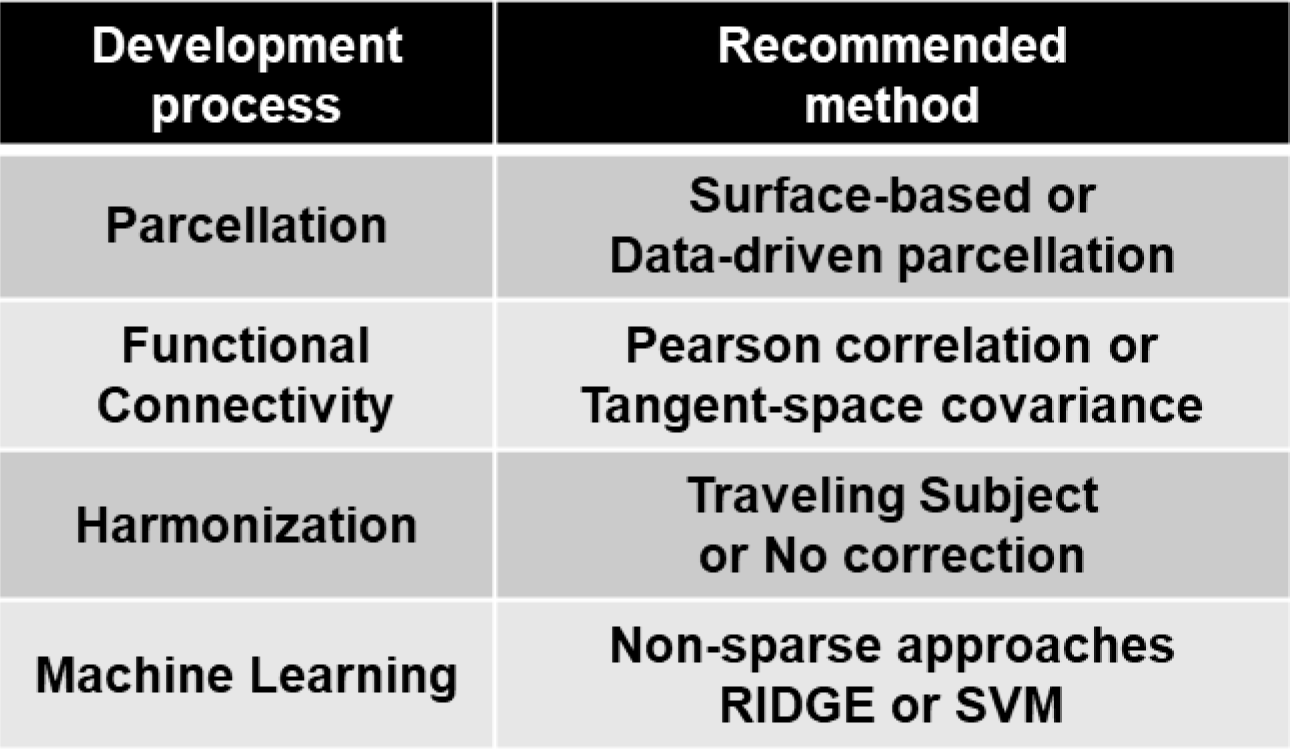
Optimal method for development of practical biomarker.

| **Development process** | **Recommended method** |
| --- | --- |
| **Parcellation** | **Surface-based or Data-driven parcellation** |
| **Functional Connectivity** | **Pearson correlation or Tangent-space covariance** |
| **Harmonization** | **Traveling Subject or No correction** |
| **Machine Learning** | **Non-sparse approaches**<br>**RIDGE or SVM** |

### 3.2. Evaluation of pipeline generality by dataset-role swapping

We identified the markers that show high diagnostic performance even for the validation dataset and the pipelines for constructing them. In view of the data dependency of machine learning, we constructed and verified markers using dataset II as a discovery dataset to check the generality of the results regarding marker performance rankings and important factors (Supplemental Figures 1 & 2A). Since dataset II, unlike dataset I, was acquired with multiple protocols, it is not just a cross validation. We constructed MDD diagnostic biomarkers using all the pipelines, except those that include traveling-subject harmonization because dataset II has no such corresponding traveling-subject dataset. We obtained AUC, MCC, sensitivity, and specificity from both datasets and calculated the averaged composite scores and instabilities. For each subject, the diagnostic probabilities were calculated by two markers that were constructed using the same pipeline before and after dataset-role swapping to evaluate the stability of the discrimination results. After comparing the averaged composite scores before and after dataset-role swapping, the effectiveness of surface-based Glasser parcellation became remarkable, although a similar tendency was observed using the datasets in the original role (Figure 4A). For FC, the effectiveness of Pearson’s full correlation and tangent-space covariance was again confirmed (Figure 4B). For the harmonization process, the effectiveness of the no-correction option compared to ComBat remained the same, although their differences became smaller (Figure 4C). There was no change in the effectiveness of the non-sparse method in the machine learning process (Figure 4D). Each pipeline was comprehensively ranked based on its averaged composite score and the instability value of the diagnostic markers constructed before and after dataset-role swapping and the correlation coefficient of the diagnostic probability (Table 5). All five indicators used for the rankings were standardized in all the pipelines, and the signs of the instability values were inverted.

**Figure 4.**
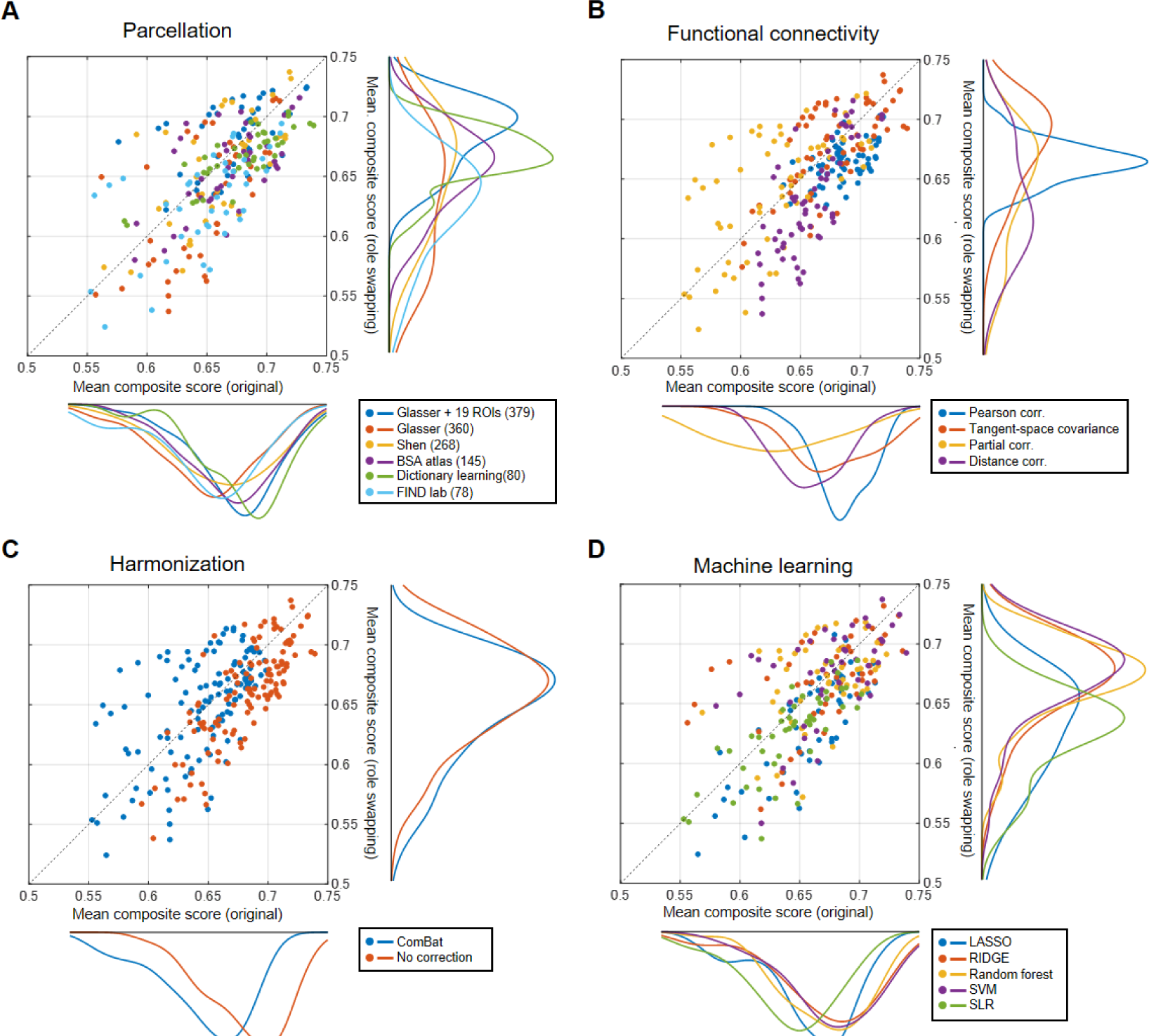
Dataset-role swapping to exclude discovery dataset-dependent pipelines. *A-D)* Distribution of averaged composite scores for each process before and after dataset-role swapping.

**Table 5.** Top 10 superior pipelines for development of MDD diagnostic biomarker: 240 constructed MDD diagnostic biomarkers were respectively ranked in descending order of averaged composite scores and in ascending order of instability. Total ranking was determined by summation of composite score ranking and instability ranking.

| Pipeline ranking | Parcellation | FC | Harmonization | Machine learning | Dataset I → II |  | Dataset II → I |  | Correlation of diagnostic prob. | Total scores |
| --- | --- | --- | --- | --- | --- | --- | --- | --- | --- | --- |
|  |  |  |  |  | Composite score | Instability | Composite score | Instability |  |  |
| #1 | Glasser + 19 ROIs (379) | Tangent | No correction | SVM | 1.825 | 1.334 | 1.640 | 0.319 | 0.761 | 5.878 |
| #2 | Glasser + 19 ROIs (379) | Tangent | No correction | RIDGE | 1.809 | 1.423 | 1.620 | 0.256 | 0.658 | 5.765 |
| #3 | Glasser + 19 ROIs (379) | Pearson | ComBat | Random forest | 0.607 | 1.289 | 1.012 | 0.982 | 1.721 | 5.611 |
| #4 | Shen (268) | Pearson | No correction | Random forest | 1.360 | 0.823 | 0.647 | 1.153 | 1.573 | 5.555 |
| #5 | Dictionary Learning (80) | Tangent | No correction | SVM | 1.972 | 0.965 | 0.898 | 1.130 | 0.359 | 5.324 |
| #6 | FIND lab (78) | Tangent | No correction | SVM | 1.281 | 0.926 | 0.936 | 1.054 | 1.086 | 5.283 |
| #7 | Dictionary learning (80) | Tangent | No correction | RIDGE | 1.893 | 1.031 | 0.936 | 1.243 | 0.176 | 5.280 |
| #8 | Glasser + 19 ROIs (379) | Pearson | No correction | Random forest | 1.087 | 0.807 | 0.754 | 1.196 | 1.265 | 5.109 |
| #9 | Dictionary learning (80) | Pearson | ComBat | Random forest | 0.686 | 1.209 | 0.277 | 1.228 | 1.535 | 4.935 |
| #10 | Glasser + 19 ROIs (379) | Pearson | No correction | SVM | 1.458 | 0.971 | 1.122 | 0.540 | 0.782 | 4.873 |

### 3.3. Evaluation of marker similarity

We investigated the similarities among diagnostic biomarkers constructed from the top 10 identified pipelines. The mean ± SEM of the composite scores in the discovery dataset for the top 10 markers was 0.724 ± 0.010 (Figure 5A). The corresponding AUC, accuracy, sensitivity, specificity, and MCC scores were 0.775 ± 0.006, 0.682 ± 0.012, 0.761 ± 0.009, 0.663 ± 0.011, and 0.345 ± 0.014 (Supplemental Table 3). On the other hand, the mean ± SEM of the composite scores in the independent validation dataset was0.710 ± 0.005. The corresponding AUC, accuracy, sensitivity, specificity, and MCC scores were 0.743 ± 0.005, 0.674 ± 0.005, 0.711 ± 0.018, 0.649 ± 0.015, and 0.355 ± 0.010. The diagnostic results of any pair of top 10 diagnostic markers showed a very high concordance rate (Figure 5B, Sorensen-Dice coefficient index: 0.791 ± 0.008, mean ± SEM). Each marker has 100 classifiers and 100 kinds of weight or importance values for each functional connectivity. The summation of the absolute weights or the importance of each marker was regarded as the degree of its contribution for classification, and we confirmed whether the top 5% contribution FCs had network usage similarity among the markers. Since each parcellation has its own ROIs for dividing the brain, it is impossible to simply compare high contribution FC patterns among markers with different parcellations. Therefore, we put the high contribution FCs into a common brain map called brain networks proposed by Yeo et al. and compared the similarity of the usage rates of the networks to which each FC belongs (Supplemental Figure 3). Since the existence probability of all the FCs on the brain network is different for each parcellation, we compared the network utilization rates between any two markers after correcting them. As a result, significant higher correlations were observed between the markers constructed by the pipelines, including all the parcellation and functional connectivity estimation patterns, except for the combination of dictionary-learning parcellation and tangent-space covariance (Figure 5C, D).

**Figure 5.**
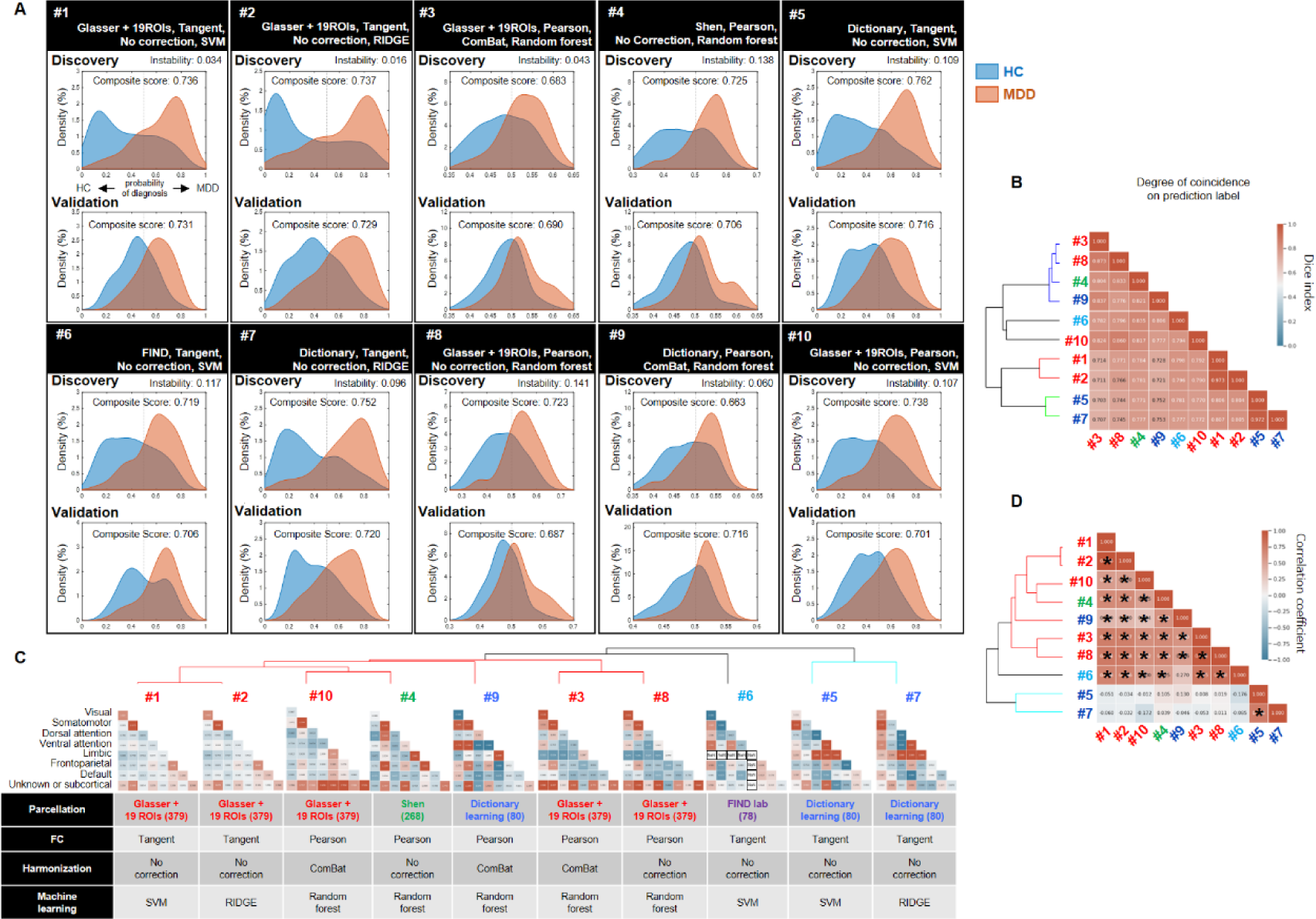
Diagnosis and weight similarity among top 10 high-performance biomarkers: *A)* Classification performance of top 10 MDD diagnostic biomarkers. Upper row shows correlation coefficient between top markers in diagnostic probability. Probability distribution for diagnosis of MDD in dataset I (discovery 10-fold CV test, upper) and dataset II (validation test, bottom). MDD and HC distributions are depicted in red and blue. *B)* Correlation coefficient between top markers in diagnostic probability. *C, D)* Similarity of important FCs’ corrected network usage amount among top 10 markers. Two-thirds combination showed a significantly higher correlation with corrected network usage amount (D, permutation test, *p<0.05).

### 3.4. Application of pipelines to other disorders

Finally, we tested whether the high-performance biomarkers of other mental disorders can be constructed with higher-ranked pipelines identified using the MDD datasets (Supplemental Figure 1) since these mental disorders also have no objective diagnostic biomarkers. Using the top 10 pipelines shown in Table 5, we verified whether diagnostic markers for ASD and SCZ can be constructed by the same procedure as that for creating diagnostic markers for MDD. In the top 10 pipelines, which did not contain a ComBat option, the diagnostic markers for ASD and SCZ were successfully constructed with equal to or higher classification performances than the diagnostic markers for MDD (Figure 6).

**Figure 6.**
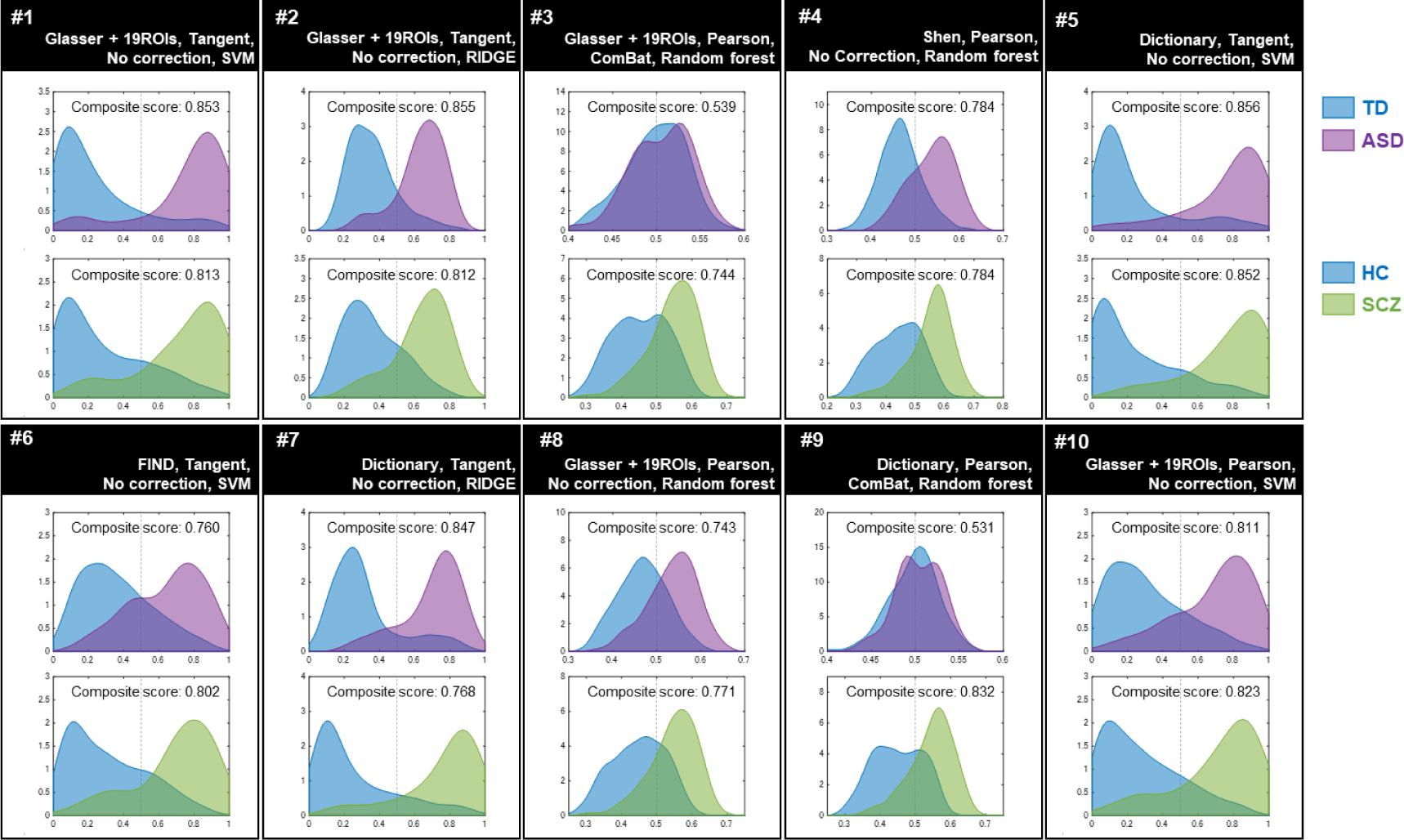
Development of ASD and SCZ diagnostic biomarkers using top 10 analysis pipelines: Constructing ASD and SCZ diagnostic biomarkers using top 10 pipelines, showing high classification and generalization performances in constructing MDD diagnostic biomarkers. Eight of top 10 markers show high classification performances on composite scores. TD or HC distributions are depicted in blue, ASD in purple, and SCZ in green.

## 4. Discussion

In this study, we conducted a comprehensive evaluation of the analysis pipelines of resting-state functional connectivity (rsFC) biomarkers using a large-scale, multi-site fMRI dataset for major depressive disorder (MDD) and healthy controls (HCs) (1162 participants from eight imaging sites). We explored option combinations in four subprocesses of the analysis pipeline: six types of brain parcellation, four types of FC computation, three types of site difference harmonization, and five types of machine learning methods. In total 360 biomarkers were constructed using the SRPBS dataset acquired with a unified protocol (713 participants from four imaging sites) as the discovery dataset and their classification performances were evaluated with the dataset from other independent projects acquired with heterogeneous protocols (449 participants from four imaging sites) as the validation dataset. We identified the best options in each of four subprocesses based on both the cross-validated classification performance within the discovery dataset and the classification performance for the validation dataset. To find the best options shared by the rsFC biomarkers constructed with a dataset acquired with a unified protocol and those constructed with a dataset with a heterogenous protocol, we repeated the same procedure after swapping the role of the two datasets. We found that the pipelines tended to result in high classification performance, including Glasser’s parcellation, tangent covariance, no harmonization and such non-sparse classifiers as ridge logistic regression and support vector machine (SVM). Then we investigated the diagnosis and weight similarity between the top 10 biomarkers and observed commonality, except in two biomarkers using both data-driven parcellation and FC computation. Finally, we applied the top 10 pipelines to the datasets of other mental disorders (autism spectrum disorder and schizophrenia) and 8 of 10 biomarkers showed sufficient classification performance. Our results support the construction of standardized pipelines for multi-site and multi-disorder biomarkers.

Regarding brain parcellation, Glasser parcellation was the most effective followed by the dictionary learning method. Glasser parcellation is surface-based. Other studies reported that surface-based parcellation, which was superior for detecting brain activity with lower signal contamination than volume-based parcellation by 2D smoothing (Brodoehl et al., 2020), increased statistical power even in lower spatial and temporal resolution cases (Anticevic et al., 2008; Coalson et al., 2018). This lower signal contamination might improve the diagnostic performance for the validation dataset, which consists of the fMRI data acquired with different protocols at various sites from the discovery dataset. The effectiveness of dictionary learning was already reported in a previous benchmarking study (Dadi et al., 2019). Although dictionary learning worked the best when it was learned with datasets acquired with the unified protocol, its performance degraded significantly when it was learned with the dataset acquired with heterogeneous protocols. Interestingly, the number of ROIs of Glasser parcellation is 379, the highest in our exploration; dictionary learning had only 80, the lowest.

Regarding FC, we confirmed that Pearson’s full correlation and tangent-space covariance were better than the other options. The latter had slightly higher performance than the former, although it was not statistically significant. Tangent-space covariance’s effectiveness was also consistent with previous reports (Dadi et al., 2019; Yang et al., 2022). Since Pearson’s full correlation uses the averaged fMRI signals within each ROI, this approach overlooks the spatial patterns of voxel-wise or vertex-wise signals within individual ROIs. To exploit the information of voxel patterns within each ROI, we tested the recently-proposed distance correlation (Yoo et al., 2019). However, the average performance was not good, indicating the high sensitivity of voxel patterns to protocol and scanner differences.

Regarding the harmonization method, we found no performance differences between the traveling-subject (TS) harmonization method and the no-correction method, although Combat’s performance degraded when the SRPBS dataset was used as discovery datasets. When we trained the biomarkers with the heterogeneous datasets where Combat is the only method for harmonization, we did not find a difference between no-correction and Combat (Supplemental Figure 2B). In summary, we did not find a clear effect of harmonization on the classification performance. One possible reason is that the pattern of disease factors is sufficiently different from the pattern due to site differences and machine learning automatically weighs disease factors (Abraham et al., 2017; Yamashita et al., 2019). However, note that harmonization remains important when closely interpreting constructed biomarkers. For example, a previous study using sparse classifiers to construct MDD biomarkers showed that the number of relevant FCs increased after TS harmonization compared with the no-correction method (Yamashita et al., 2020).

For the machine learning method, our results showed that non-sparse machine learning methods were preferred, a finding that is consistent with a previous report (Dadi et al., 2019). This result implies that the functional connectivity required for MDD diagnoses is spread throughout the brain. Although the non-sparse methods worked better as long as the classification performance was considered, sparse classifiers, which can select a small number of important FCs (Yamashita et al., 2020), are advantageous if applications target such specific FCs as FC neurofeedback (Megumi et al., 2015; Yamashita et al., 2017) and the identification of drug discovery target brain areas (Ichikawa et al., 2020).

The similarity analysis of the diagnosis and FC weights of the top 10 pipelines showed that diagnosis patterns were highly similar among the 10 markers and their weight patterns were highly similar among the eight biomarkers. Two markers, which showed a low weight similarity to the remaining markers, used a combination of dictionary-learning parcellation and tangent-space covariance, both of which are data-driven methods. With multiple data-driven methods, we can discriminate the MDD characteristics that are different from markers using the other pipelines. An interesting future study might take advantage of the different characteristics of multiple biomarkers using the ensemble learning framework to provide more robust discrimination results.

Although this study carried out a search for the optimal diagnostic markers of construction pipelines by developing MDD diagnostic biomarkers, other mental disorders also require the establishment of objective biomarkers. We showed that eight of the top 10 pipelines might also be effective in developing diagnostic markers for ASD and SCZ in addition to MDD. Since diagnostic markers have been constructed for these three mental disorders, each subject can obtain in a single fMRI acquisition the diagnostic probability for each disorder from the three diagnostic biomarkers. Combining multiple diagnostic probabilities based on biophysiological backgrounds is expected to fuel patient classifications regardless of conventional categorical diagnoses. We will also attempt to study patient-clustering methods using a multiple disorder dataset, an attempt also consistent with the RDoC concept, which seeks precision medicine for psychiatry (Insel, 2014). Regarding the low diagnostic performance of the two ASD diagnostic biomarkers, both were constructed using pipelines, including a combination of Pearson’s full correlation, ComBat harmonization, and the random forest method into a discovery dataset. Since the two discovery dataset sites consisted of only typical development subjects and ASD patients were concentrated in one site, excessive correction was caused by ComBat, and the characteristics of the ASD patients on FC seemed lost. In fact, if diagnostic markers are created in a pipeline containing ComBat for only two sites including ASD, their diagnostic performance is improved (Supplemental Figure 4).

In summary, we searched for a pipeline with excellent generalization performance and excellent marker construction from a combination search of comprehensive methods based on fMRI data acquired at multiple large-scale facilities and obtained multiple candidates. Our identified pipeline is likely to be applicable to other psychiatric disorders, and we expect that a combination of multiple fMRI markers will support new understanding of such disorders. We hope that the widespread use of fMRI markers in clinical practice will shorten the treatment period for patients and lead to the development of new treatment methods for those for whom no effective treatment method currently exists.

## Limitations

In this study, although the fMRI data were acquired from multiple large-scale facilities, they were only in Japan. In our previous study (Yamashita et al., 2020), we applied our MDD biomarker to an fMRI dataset shared by OpenNeuro as an independent validation dataset from a foreign country. However, since half of its fMRI data were inexplicably replaced after the previous publication by the providers, we could not obtain sufficient performance when we applied any of the markers constructed in this study to the dataset of our latest version (Supplemental Figure 5). Although we carefully investigated the replaced data to identify the cause, we failed. This fact suggests the urgency of data acquisition under internationally-controlled protocols and data sharing in a common format for investigating the cause and development of more practical markers. In addition, regarding constructing autism spectrum disorder and schizophrenia diagnostic markers, we haven’t yet conducted a verification test using external validation data.

## Data availability

We are a registered member of the Decoded Neurofeedback (DecNef) Project Brain Data Repository (https://bicr.atr.jp/decnefpro/data), and the data for this study are available from its website on reasonable request by qualified researchers. See Tanaka et al., 2021 for more detailed information about the datasets.

