## Supplemental File for "Comprehensive evaluation of pipelines for diagnostic biomarkers of major depressive disorder using multi-site resting-state fMRI datasets"

Supplemental Table 1. Imaging protocols for resting-state fMRI in dataset I & II

| **Dataset** | **Dataset I** | | | | **Dataset II** | | | |
|---|---|---|---|---|---|---|---|---|
| **Site** | **Center of Innovation in Hiroshima university** | **Kyoto university** | **Showa university** | **University of Tokyo** | **Hiroshima Kajikawa hospital** | **Hiroshima rehabilitation center** | **Hiroshima university hospital** | **Yamaguchi university** |
| Abbreviation | COI | KUT | SWA | UTO | HKH | HRC | HUH | UYA |
| MRI scanner | *Siemens Verio* | *Siemens TimTrio* | *Siemens Verio* | *GE MR750w* | *Siemens Spectra* | *GE Signa HDxt* | *GE Signa HDxt* | *Siemens Skyra* |
| Magnetic field strength | 3.0 T | | | | 3.0 T | | | |
| Channels per coil | 12 | 32 | 12 | 24 | 12 | 8 | 8 | 20 |
| Field-of-view (mm) | 212 × 212 | | | | 192 × 192 | 256 × 256 | 256 × 256 | 220 × 220 |
| Matrix | 64 × 64 | | | | | | | |
| Number of slices | 40 | | | | 38 | 32 | 32 | 34 |
| Number of volumes | 240 | | | | 107 | 143 | 143 | 200 |
| In-plane resolution (mm) | 3.3125 × 3.3125 | | | | 3.0 × 3.0 | 4.0 × 4.0 | 4.0 × 4.0 | 3.4 × 3.4 |
| Slice thickness (mm) | 3.2 | | | | 3 | 4 | 4 | 4 |
| Slice gap (mm) | 0.8 | | | | 0 | 0 | 0 | 1 |
| TR (ms) | 2500 | | | | 2,700 | 2,000 | 2,000 | 2,500 |
| TE (ms) | 30 | | | | 31 | 27 | 27 | 30 |
| Total scan time (min:s) | 10:00 | | | | 5:00 | 4:46 | 5:00 | 8:28 |
| Flip angle (degree) | 80 | | | | 90 | 90 | 90 | 80 |
| Slice acquisition order | Ascending | | | | Ascending | Ascending (interleaved) | Ascending (interleaved) | Ascending |
| Phase encoding | AP | PA | PA | PA | AP | AP | PA | PA |
| Eyes closed/ Open/ fixated | Fixated | | | | Fixated | Fixated | Fixated | Closed |

Supplemental Table 2. Imaging protocols for resting-state fMRI in traveling-subject-harmonization dataset (Yamashita et al., PLoS Biol, 2019)

| Site | ATR *Tim Trio* | ATR *Verio* | Center of Innovation in Hiroshima university | Hiroshima University Hospital | Hiroshima Kajikawa hospital | Kyoto Prefectural university of medicine | Showa university | Kyoto university Tim Trio | University of Tokyo |
|---|---|---|---|---|---|---|---|---|---|
| Abbreviation | ATT | ATV | COI | HUH | HKH | KPM | SWA | KUT | UTO |
| MRI scanner | Simens | | | GE | Simens | Philips | Simens | | GE |
| | *Tim Trio* | *Verio* | | *Signa HDxt* | *Spectra* | *Achieva* | *Verio* | *Tim Trio* | *MR750W* |
| Magnetic field strength | 3.0T | | | | | | | | |
| Channels per coil | 12 | | | 8 | 12 | 8 | 12 | 32 | 24 |
| Field-of-view (mm) | 212 × 212 | | | 256 × 256 | 192 × 192 | | 212 × 212 | | |
| Matrix | 64 × 64 | | | | | | | | |
| Number of Slices | 40 or 39 | 39 | 40 | 32 | 38 | 39 | 40 | | |
| Number of volumes | 240 | | | 143 | 107 | 194 | 240 | | |
| In-plane resolution (mm) | 3.3125 × 3.3125 | | | 4.0 × 4.0 | 3.0 × 3.0 | | 3.3125 × 3.3125 | | |
| Slice thickness (mm) | 3.2 | | | 4 | 3 | | 3.2 | | |
| Slice gap (mm) | 0.8 | | | 0 | | | 0.8 | | |
| TR (ms) | 2,500 | | | 2,000 | 2,700 | 2,000 | 2,500 | | |
| TE (ms) | 30 | | | 27 | 31 | 30 | | | |
| Total scan time (min:s) | 10:00 | | | 5:00 | 5:00 | 6:30 | 10:00 | | |
| Flip angle (degree) | 80 | | | 90 | | 80 | | | |
| Slice acquisition order | Ascending | | | Ascending (interleaved) | Ascending | | | | |
| Phase encoding | PA | | AP | PA | AP | | PA | | |
| Eyes closed/ Open/ Fixated | Fixated | | | | | Closed | Fixated | | |

Supplemental Table 3. Diagnostic performances both in discovery and validation datasets

| Pipeline ranking | Parcellation | FC | Harmonization | Machine learning | Discovery dataset (dataset I) 10-fold CV test | | | | | Validation dataset (dataset II) | | | | |
|---|---|---|---|---|---|---|---|---|---|---|---|---|---|---|
| | | | | | AUC | Accuracy | Sensitivity | Specificity | MCC | AUC | Accuracy | Sensitivity | Specificity | MCC |
| #1 | Glasser + 19 ROIs (379) | Tangent | No correction | SVM | 0.798 | 0.694 | 0.746 | 0.681 | 0.349 | 0.761 | 0.693 | 0.758 | 0.649 | 0.400 |
| #2 | Glasser + 19 ROIs (379) | Tangent | No correction | RIDGE | 0.798 | 0.704 | 0.725 | 0.699 | 0.349 | 0.761 | 0.698 | 0.713 | 0.687 | 0.394 |
| #3 | Glasser + 19 ROIs (379) | Pearson | ComBat | Random forest | 0.721 | 0.643 | 0.739 | 0.618 | 0.288 | 0.729 | 0.645 | 0.697 | 0.610 | 0.301 |
| #4 | Shen (268) | Pearson | No correction | Random forest | 0.777 | 0.668 | 0.790 | 0.637 | 0.344 | 0.742 | 0.675 | 0.661 | 0.685 | 0.341 |
| #5 | Dictionary Learning (80) | Tangent | No correction | SVM | 0.821 | 0.720 | 0.797 | 0.701 | 0.408 | 0.752 | 0.682 | 0.691 | 0.676 | 0.361 |
| #6 | FIND lab (78) | Tangent | No correction | SVM | 0.765 | 0.679 | 0.768 | 0.657 | 0.345 | 0.726 | 0.661 | 0.820 | 0.552 | 0.373 |
| #7 | Dictionary Learning (80) | Tangent | No correction | RIDGE | 0.815 | 0.706 | 0.775 | 0.688 | 0.379 | 0.758 | 0.686 | 0.680 | 0.691 | 0.366 |
| #8 | Glasser + 19 ROIs (379) | Pearson | No correction | Random forest | 0.770 | 0.684 | 0.768 | 0.662 | 0.350 | 0.718 | 0.664 | 0.629 | 0.687 | 0.313 |
| #9 | Dictionary Learning (80) | Pearson | ComBat | Random forest | 0.700 | 0.625 | 0.710 | 0.604 | 0.253 | 0.747 | 0.670 | 0.770 | 0.602 | 0.367 |
| #10 | Glasser + 19 ROIs (379) | Pearson | No correction | SVM | 0.785 | 0.701 | 0.790 | 0.679 | 0.382 | 0.735 | 0.666 | 0.685 | 0.653 | 0.332 |
| Mean ± SEM | | | | | 0.775 ± 0.012 | 0.682 ± 0.009 | 0.761 ± 0.009 | 0.663 ± 0.011 | 0.345 ± 0.014 | 0.743 ± 0.005 | 0.674 ± 0.005 | 0.710 ± 0.018 | 0.649 ± 0.015 | 0.355 ± 0.010 |

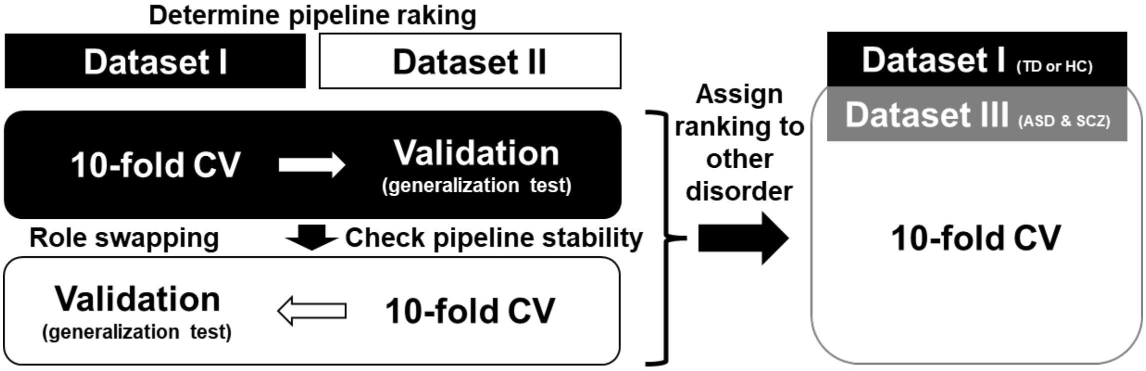


**Supplemental Figure 1. Schema of our study:** Pipeline ranking was constructed using datasets I and II composed of depressed and healthy subjects. Using top 10 pipeline determined by constructing MDD diagnostic markers, we constructed diagnostic markers for ASDs and SCZs. Data for ASD and SCZ were included in dataset 3 and were used together with data for TD and HC from dataset 1.

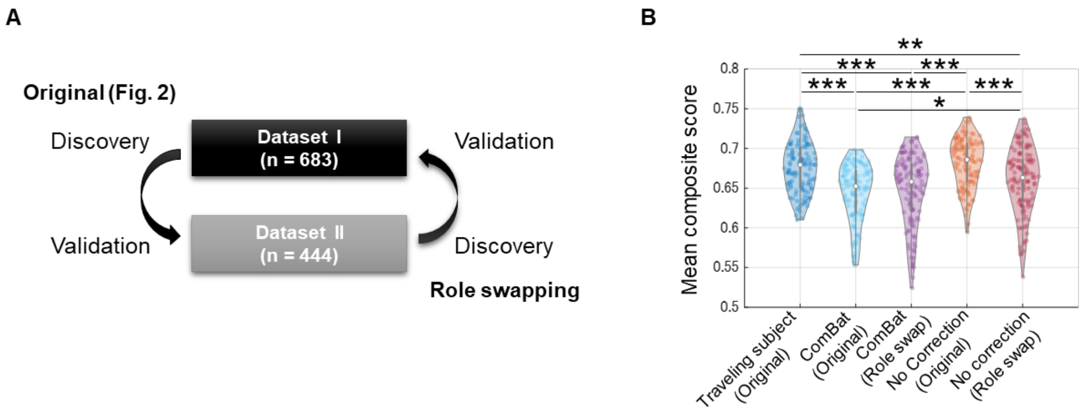


**Supplemental Figure 2. Dataset-role swapping:** *A)* Dataset II was used as discovery dataset and dataset I was used for validation. *B)* Role swapping significantly reduced diagnostic performance of markers constructed using pipelines selected "no correction."

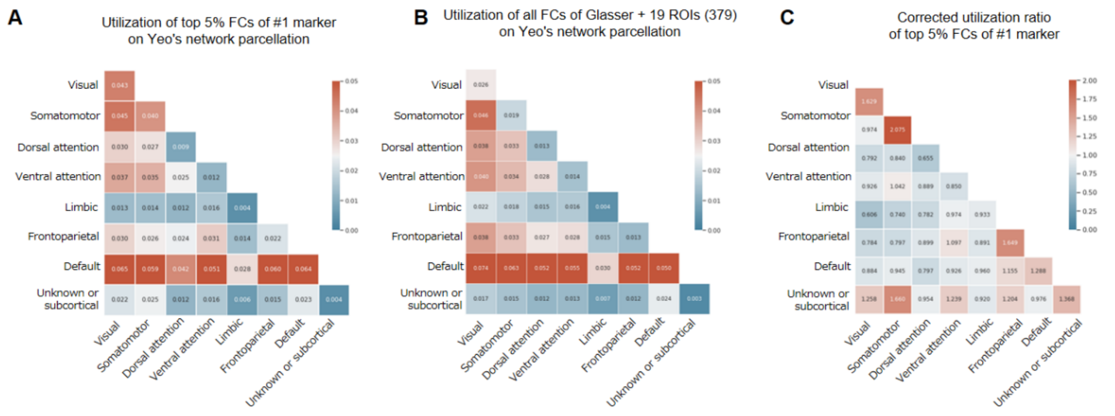


**Supplemental Figure 3. Procedure comparing important FCs between each diagnostic marker after applying identical Yeo's network parcellation:** *A)* Existence probability is calculated by classifying all FCs included in selected parcellation using Yeo's brain network. Case of Glasser + 19 ROIs (379) is displayed. *B)* In marker of #1 using Glasser + 19 ROIs (379) parcellation, top 5% FCs of absolute values of weight are extracted and applied to Yeo's network to calculate usage ratio, although bias of usage ratio of parcellation itself is ignored at this stage. C*)* By dividing usage ratio obtained in *B)* by parcellation-specific usage ratio in *A)*, corrected network usage amount, which can be compared between markers, is calculated.

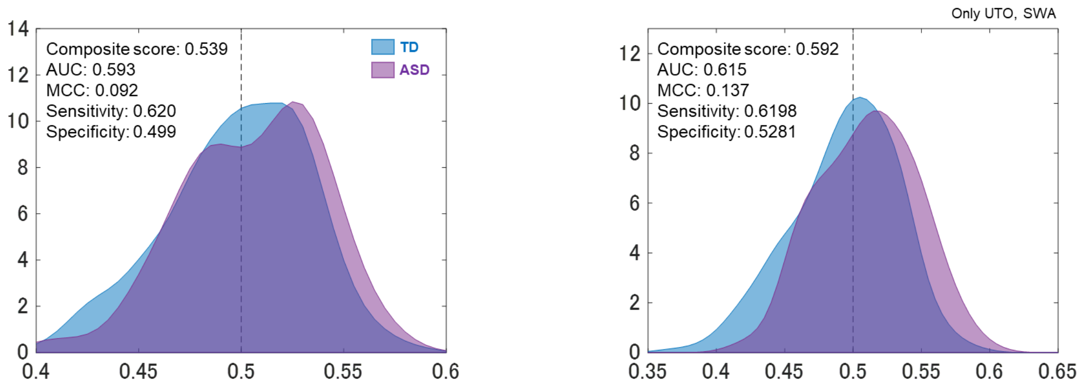


**Supplemental Figure 4. Impact on ComBat of including sites consisting only of healthy people**

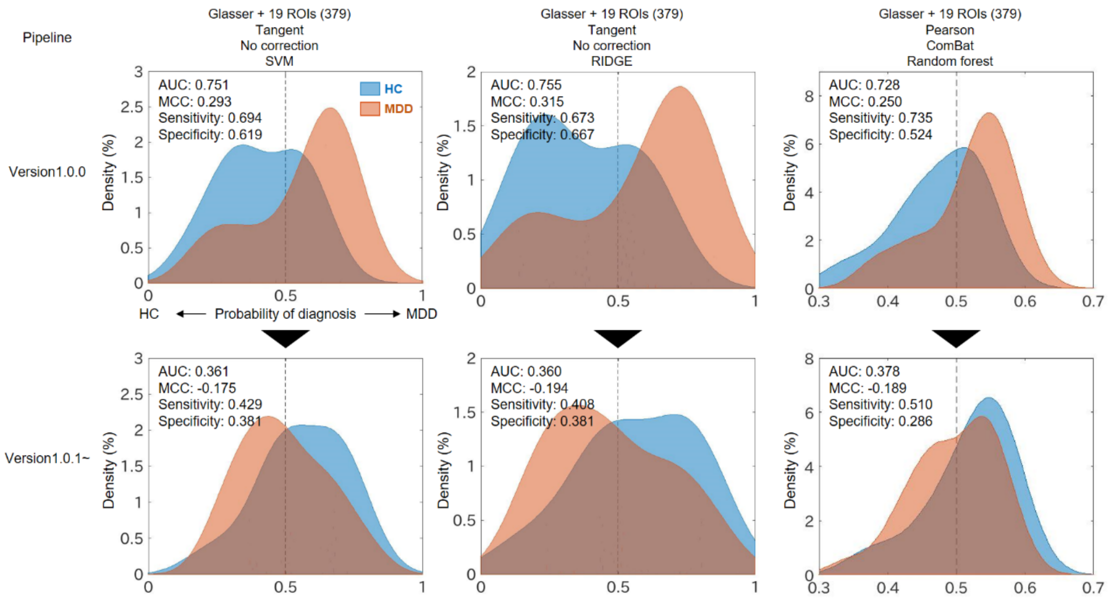


**Supplemental Figure 5. Application top three markers to foreign MDD dataset:**

*top)* Results of applying markers constructed by top three pipelines to same dataset (https://openneuro.org/datasets/ds002748/versions/1.0.0) as previous study. *bottom)* Results of applying them to dataset of latest version (https://openneuro.org/datasets/ds002748/versions/1.0.5).
